# Structural evolutionary analysis predicts functional sites in the artemisinin resistance malaria protein K13

**DOI:** 10.1101/346668

**Authors:** Romain Coppée, Daniel C. Jeffares, Audrey Sabbagh, Jérôme Clain

**Affiliations:** UMR 216 MERIT, Institut de Recherche pour le Développement, Université Paris Descartes, Sorbonne Paris Cité, Paris, France; Department of Biology, University of York, Wentworth Way, York, YO10 5DD, UK; Centre National de Référence du Paludisme, Hôpital Bichat-Claude Bernard, Assistance Publique des Hôpitaux de Paris, France

## Abstract

K13 is an essential Plasmodium falciparum protein that plays a key role in malaria resistance to artemisinins. Although K13 resembles BTB- and Kelch/propeller-containing proteins involved in ubiquitin ligase complexes, its functional sites remain uncharacterized. Using evolutionary and structural information, we searched for the most conserved K13 sites across Apicomplexa species evolution to identify sub-regions of K13 that are likely functional. An amino acid electropositive ‘patch’ in the K13 propeller domain has a dense concentration of extraordinarily conserved positions located at a shallow pocket, suggesting a role as binding surface. When applied to experimentally-characterized BTB-Kelch proteins, our strategy successfully identifies the validated substrate-binding residues within their own propeller shallow pocket. Another patch of slowly evolving sites is identified in the K13 BTB domain which partially overlaps the surface that binds to Cullin proteins in BTB-Cullin complexes. We provide candidate binding sites in K13 propeller and BTB domains for functional follow-up studies.

## Introduction

Current efforts to control malaria are threatened by the spread in Southeast Asia (SEA) of *Plasmodium falciparum* parasites that are resistant to artemisinin derivatives (ARTs)^1^. Treatment failures are now reported in some geographic areas of SEA for the current front-line ART-based combination therapies^1–3^. In Africa, ART resistance (ART-R) is not yet established^4,5^, although some *P. falciparum* parasites from Uganda and Equatorial Guinea exhibiting high survival rate have been described^6,7^.

In parasites from SEA, ART-R is primarily conferred by single non-synonymous mutations in the *P. falciparum k13* (*pfk13*) gene^8,9^. Multiple *pfk13* ART-R mutations have emerged concomitantly in the early 2000’s until a specific, multidrug-resistant lineage carrying the C580Y mutation became the most common, especially in the East Thailand-Cambodia-Lao PDR-Vietnam region^4,8,10–12^. The ART-R phenotype is defined as parasites exhibiting *in vivo* a delayed clearance time following an ART-based treatment^1^ and *in vitro* an increased survival rate following a brief exposure to a high dose of ART^13^.

The *pfk13* gene encodes a 726 amino acid protein (PfK13) which is essential at least during the intraerythrocytic parasite blood stage^14,15^. Both the gene ontology annotation of *pfk13* and the study of ART-resistant parasites carrying a *pfk13* mutation suggest that this protein has some regulatory functions at the protein level^16,17^. At the cell level, *pfk13* mutant parasites decelerate their development during the early intraerythrocytic stage^13,18^. At the molecular level, they exhibit an increased expression of unfolded protein response pathways^19^, lower levels of ubiquitinated proteins^17,18^, and phosphorylation of the parasite eukaryotic initiation factor-2α (eIF2α) which correlates with ART-induced latency^20^. There are some indications of the interactors with PfK13 that partially clarify its function. For example, PfK13 was immunoprecipitated with the phosphatidylinositol 3-kinase (PI3K)^17^. Ubiquitination of PI3K is also decreased in *pfk13* C580Y mutants, resulting in wide phosphatidylinositol 3-phosphate-related cellular alterations^17,21^ which roles in ART-R are still to be characterized.

PfK13 is related to the BTB-Kelch structural subgroup of Kelch-repeat proteins^8,22,23^. It possesses a BTB domain (Broad-complex, tramtrack and bric-à-brac; also known as BTB/POZ; amino acids 350-437) and a C-terminal propeller domain (also known as Kelch domain; amino acids 443-726) composed of six repeated Kelch motifs (PDB code 4YY8, unpublished). However, PfK13 exhibits specific features when compared to typical BTB-Kelch proteins, such as a poorly conserved *Apicomplexa*-specific N-terminal region, a coiled-coil-containing (CCC) domain (amino acids 212-341) located upstream of BTB, and an absence of BACK domain (for BTB And C-terminal Kelch) often found between BTB and Kelch domains^23–25^. Both BTB and propeller are protein domains known to carry binding functions. Nearly all *pfk13* mutations associated with ART-R – including the C580Y mutation – cluster in the propeller domain^8,11^, suggesting a major functional role of this domain.

Many proteins harboring a BTB domain are found in multi-subunit Cullin-RING E3 ligase complexes in which a substrate protein will be ubiquitinated and then degraded by the proteasome^24,26–28^. In those complexes, BTB mediates varying oligomerization architectures and also contributes to Cullin recruitment. The propeller domain often serves as the substrate protein receptor in BTB-Kelch and other proteins^23,24^. It usually comprises four to six repeated β-stranded Kelch motifs – also named blades – arranged in a circular symmetry (the six blades in PfK13 are named I to VI). The loops protruding at the bottom face of propeller form a shallow pocket involved in the binding of the substrate protein subsequently ubiquitinated and targeted for degradation^24^. For example, the propeller shallow pocket of BTB-Kelch KEAP1 and KLHL3 directly binds to the transcription factor Nrf2 and the kinase WNK, respectively, and controls their ubiquitination^29,30^. PfK13 may exhibit similar functions, however, its binding regions and functionally important sites remain poorly characterized.

Here, we hypothesize that functionally important sites of K13 have evolved under stronger purifying selection and could be identified by the analysis of *k13* molecular evolution across 43 *Apicomplexa* species. To examine this, we inferred substitution rates for each amino acid site of the K13 sequence, taking into account the *k13* phylogeny and the spatial correlation of site-specific substitution rates in the protein tertiary structure when possible. We identified a major functional patch of slowly evolving sites located at the bottom face of K13 propeller that form part of the shallow pocket. To show the relevance of our approach, we applied it to the propeller domain of four well-characterized BTB-Kelch proteins (KEAP1, KLHL2, KLHL3 and KLHL12) and successfully identified the functionally and structurally validated substrate-binding residues located in the pocket of these propeller domains. Another patch of slowly evolving sites was also identified in the BTB domain of K13 which partially overlaps with the surface that binds to Cullin proteins in known BTB-Cullin complexes^27^. Altogether, these findings support a crucial role of K13 in binding partner molecules and predict specific binding sites.

## Results

### K13 sequence sampling, multiple alignment and phylogenetic reconstruction

Forty-three complete amino acid sequences from distinct *Apicomplexa* species were unambiguously identified as orthologous to PfK13 in sequence databases, encompassing 21 *Plasmodium* and 22 other *Apicomplexa* K13 sequences (*Cryptosporidia*, n = 7; *Piroplasmida*, n = 7; and *Coccidia*, n = 8). The length of K13 protein sequences ranged from 506 (*Eimeria brunetti*) to 820 (*Hammondia hammondi*) amino acids (Supplementary Table 1). By visual inspection, the three annotated domains of K13 (CCC, BTB and propeller) were conserved (see the K13 sequence alignment in Supplementary Fig. 1), whereas the N-terminal region preceding the CCC domain appeared much more variable among sequences, even being absent in some of them. Since K13 sequences aligned poorly in that region, the first 234 amino acid positions of the K13 multiple alignment were removed, along with other positions showing a high divergence level and/or a gap enrichment among sequences (32 amino acid positions; Supplementary Fig. 1). The final K13 multiple sequence alignment contained 514 amino acid positions which covered the whole CCC, BTB and propeller domains. The average pairwise sequence identity in that cleaned K13 sequence alignment was 64.6%, ranging from 48.6% for the *Babesia bigemina-Cryptosporidium ubiquitum* comparison to 99.2% for the *P. chabaudi* spp. pair.

The maximum-likelihood phylogenetic tree built from the curated alignment of the corresponding *Apicomplexa k13* cDNA sequences revealed four monophyletic groups: *Cryptosporidia, Plasmodium, Piroplasmida* and *Coccidia,* all supported by high bootstrap values (≥ 98%; Supplementary Fig. 2). The group of *Hematozoa k13* sequences appeared as paraphyletic (bootstrap value = 100%), with *Piroplasmida* unexpectedly clustering with *Coccidia* (Supplementary Fig. 2). The phylogenetic relationships of *Plasmodium k13* sequences were largely consistent with the acknowledged phylogeny of *Plasmodium* species, except for bird-infecting *Plasmodium k13* sequences (*P. gallinaceum* and *P. relictum*), which appeared related to human-infecting *P. ovale* spp. sequences, although this grouping was poorly supported (bootstrap value = 47%; Supplementary Fig. 2).

### The *k13* sequence has evolved under strong purifying selection

To evaluate the selective pressure acting on *k13*, we used codon substitution models to estimate the rate of non-synonymous to synonymous substitutions, *ω* = *d*_N_/*d*_S_, across codon sites of the *k13* sequence (site models) and branches of the *k13* phylogeny (branch models). A series of nested likelihood ratio tests (LRTs) using different sets of site and branch models were carried out using the codeml tool from the PAML package^31^. When applied to the *k13* codon alignment, LRTs of codon and branch models indicated varying substitution rates *ω* both among codon sites of the *k13* sequence (M0-M3 comparison, *p* = 3.3 × 10^−225^) and among branches of the *k13* phylogeny (M0-FR, *p* = 1.9 × 10^−53^; Table 1 and Supplementary Table 2). This suggests that *k13* has evolved under a variable selective regime both across codon sites and lineages. No evidence of positive selection was found in any of the tree branches (Supplementary Fig. 3). Similarly, site models incorporating positive selection (M2a and M8) provided no better fit to the data than those including only purifying selection and neutral evolution (M1a and M7: 2ΔlnL = 0 in both cases; Table 1), thus supporting an absence of detectable adaptive selection events at any *k13* codon site over the long time scale of *Apicomplexa* evolution. Altogether, the data indicate that much of the K13 protein, except the N-terminal region, has been strongly conserved over evolutionary time.

**Table 1.**
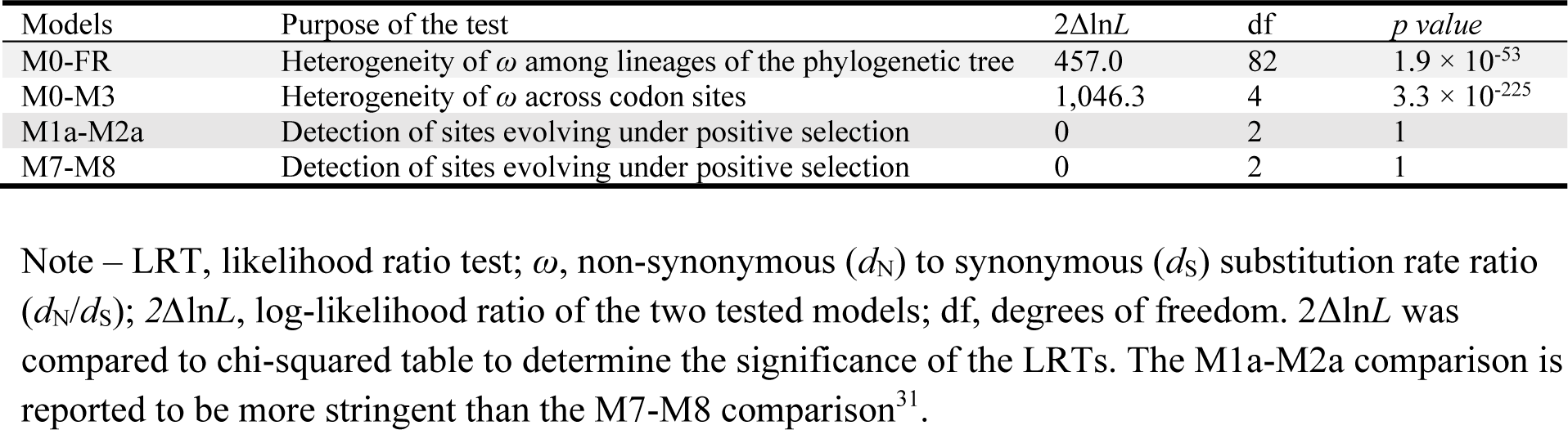
Lack of detectable positive selection in the *k13* gene using codon substitution models

When considering the one-ratio PAML values of 3,256 protein-coding genes previously estimated by Jeffares and colleagues using six *Plasmodium* species^32^, *k13* ranked among the 5% most conserved protein-coding genes of the *Plasmodium* proteome (rank: 140/3,256; Supplementary Fig. 4a). Since a significant correlation between protein length and one-ratio PAML value was evidenced in the whole dataset (Spearman’s rank correlation: *p* = 9.2 × 10^−82^, r = 0.33; Supplementary Fig. 4b), we repeated the analysis by considering only those protein sequences whose length was included in an interval of ± 100 amino acids centered on the PfK13 protein length (Spearman’s rank correlation: *p* = 0.83, r = 0.01). Again, *k13* ranked among the most conserved protein-coding genes of the *Plasmodium* proteome (sized-rank: 6/393), whereas four other five-or six-bladed Kelch protein-coding sequences showed much less intense levels of conservation than K13 (Fig. 1).

**Fig 1.**
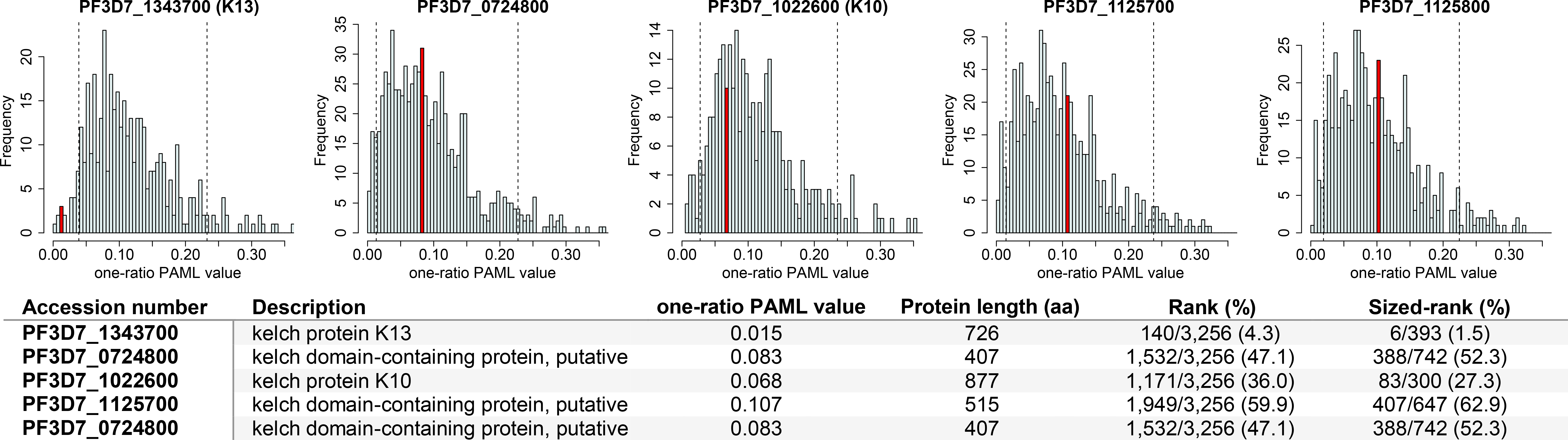
Conservation level of k13 and four other Kelch-containing protein-coding sequences compared to the all *Plasmodium* protein-coding genes. The conservation level (*d*_N_/*d*_S_) was estimated for 3,256 orthologous genes among six *Plasmodium* species (*P. falciparum*, *P. berghei*, *P. chabaudi*, *P. vivax*, *P. yoelii* and *P. knowlesi*) under the one-ratio PAML model by Jeffares and colleagues^32^. Histograms show the distribution of one-ratio PAML values for protein-coding genes whose length is comprised in an interval of ± 100 amino acids centered on each Kelch-containing protein length (sized-rank). PlasmoDB accession numbers of the five Kelch-containing proteins investigated are provided above each histogram. Red bars indicate the position of each of these Kelch-containing proteins in their respective *d*_N_/*d*_S_ distribution, with vertical dashed lines showing the five percent cutoff of the most and less conserved protein-coding genes. The table, shown below the histograms, provides the precise rank and sized-rank of each Kelch-containing protein-coding sequence. This latter corrects for a correlation between protein length and one-ratio PAML value.

### Variable levels of amino acid conservation between the annotated domains of K13

We next compared the conservation level between the annotated domains of K13 (CCC, BTB and propeller) using *ω* estimates obtained under the best fitted PAML model that indicates a variable selective regime among sites (model M3; Supplementary Table 2). First, we noted that the three domains have evolved under strong purifying selection with most sites being highly conserved during evolution (Fig. 2). BTB was however found to evolve under more intense purifying selection than either CCC (*p* = 1.6 × 10^−4^, Mann-Whitney *U* test) or propeller (*p* = 1.0 × 10^−3^, Mann-Whitney *U* test), but no difference in *ω* estimates was detected between CCC and propeller (*p* = 0.75, Mann-Whitney *U* test; Fig. 2 and Supplementary Table 3). To confirm these results, we inferred the site-specific substitution rate at the protein level using the FuncPatch server which takes into account the spatial correlation of site-specific substitution rates in the protein tertiary structure^33^ (hereafter called *λ* substitution rate). *λ* could not be inferred for the CCC domain (because of the lack of a resolved 3D structure), but the analysis confirmed that BTB was more conserved than propeller over evolutionary time (*p* = 5.4 × 10^−5^, Mann Whitney *U* test; Supplementary Table 4).

**Fig 2.**
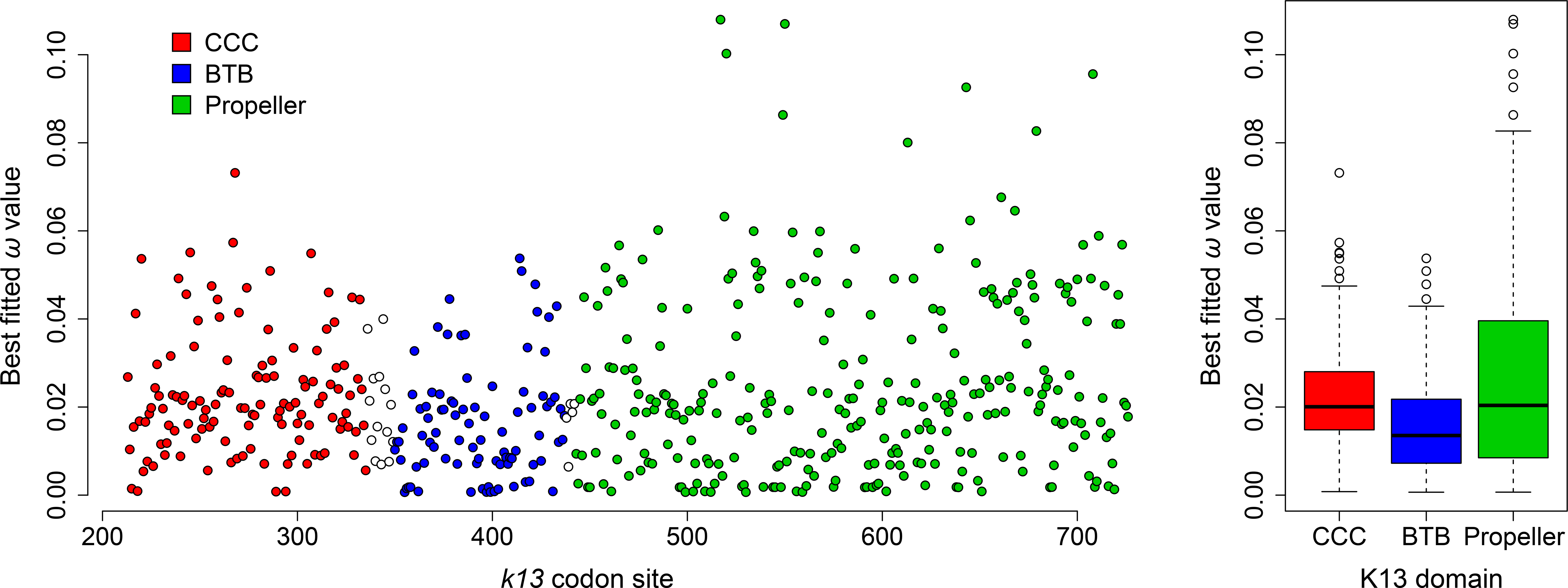
Conservation level between the three annotated domains of K13. The scatter plot (*left*) shows the conservation level of *k13* codon sites using the *pfk13* sequence as reference for the site numbering(starting at codon 213). White circles correspond to inter-domain positions. All codon sites were reported to evolve under strong purifying selection, with *ω* drastically < 1. The box-whisker plot (*right*) shows that BTB evolves under more intense purifying selection compared to either CCC or propeller, using non-parametric Mann-Whitney *U* test (*p* < 0.05). Box boundaries represent the first and third quartiles and the length of whiskers correspond to 1.5 times the interquartile range. We used the *ω* estimates obtained under the best fitted PAML model that indicates a variable selective regime among sites.

### The BTB domain of K13 resembles that of KCTD proteins and exhibits a predicted functional patch

We then performed a more extensive study of the BTB and propeller domains of K13 because of their likely role in mediating K13 functions and the availability of their tertiary structures. To detect patches of slowly evolving amino acid sites in the BTB-propeller structure, we focused on the site-specific, spatially correlated, substitution rate *λ* at the amino acid level. This rate has been shown to provide a more reliable estimation of the conservation level at amino acid sites compared to standard substitution estimates, especially in the case of highly conserved proteins^33,34^.

Although K13 is related to the BTB-Kelch structural subgroup of proteins^8^, the BTB domain of K13 exhibits atypical features compared to BTB-Kelch proteins. First, the K13 BTB fold appeared shortened, lacking the A6 helix and the N-and C-terminal extensions^27^, similar to Elongin C (Fig. 3a). Second, the primary sequence of K13 BTB grouped with those of the KCTD protein family rather than of other BTB-containing protein families (Fig. 3b). Finally, K13 BTB exhibited a higher similarity in tertiary structure with the BTB domain of KCTD17 compared to those of Elongin C and KEAP1: the root-mean-square deviations (RMSDs) of atomic positions for BTB domains of K13-KCTD17, K13-Elongin C and K13-KEAP1 were 1.13 Ångström (Å), 2.33 Å and 2.17 Å, respectively.

**Fig 3.**
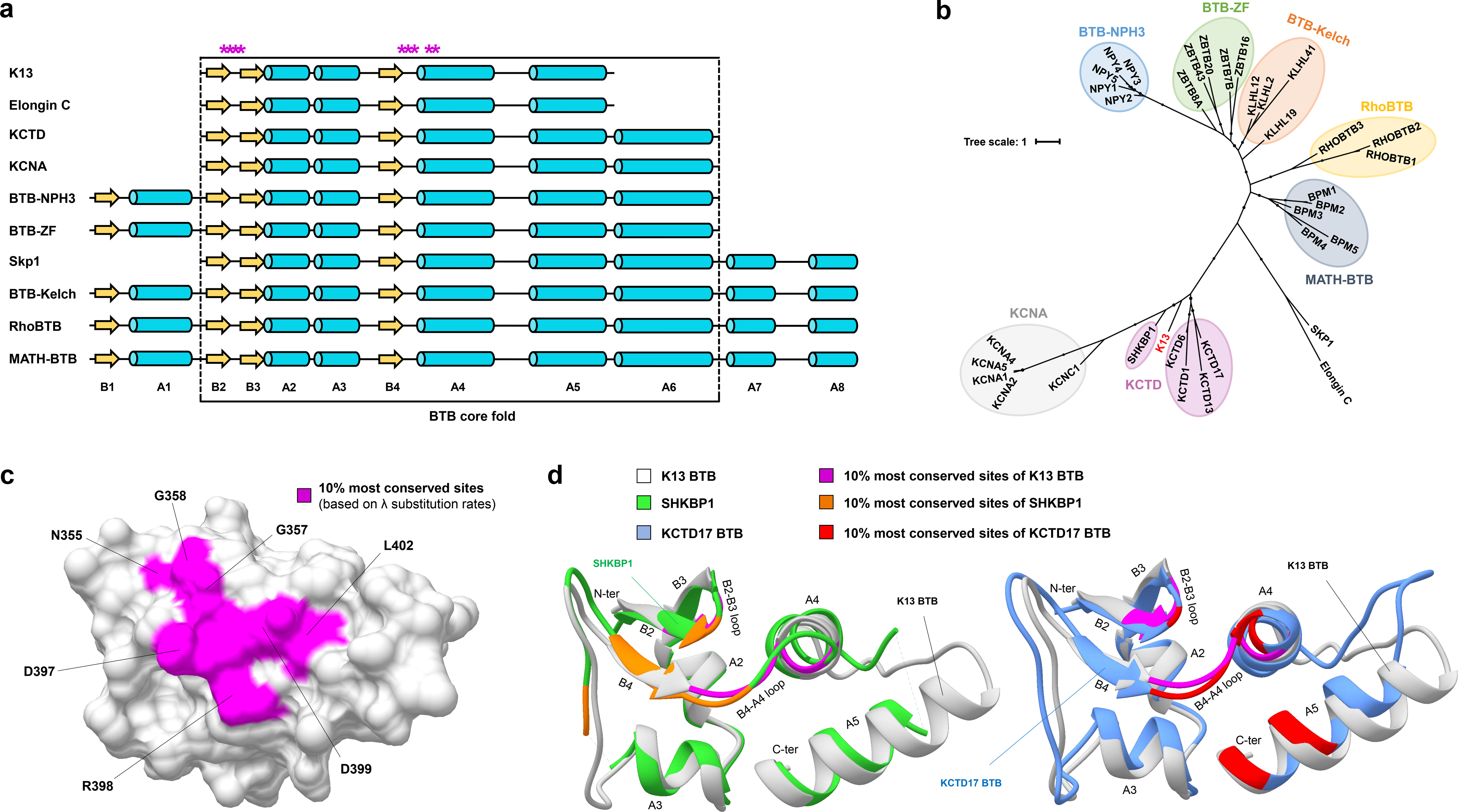
Conservation and structure homology of the K13 BTB fold. **a.** Linear schematic representation of the BTB fold of some BTB-containing protein families. Yellow arrows and cyan cylinders represent strands and helices, respectively. Stars in magenta correspond to the 10% most conserved K13 BTB amino acid sites (based on the *λ* substitution rate ranking, FuncPatch analysis, Supplementary Data 1). Structural elements are labelled. **b.** Unrooted phylogenetic tree of the BTB protein superfamily using a few reference sequences per BTB-containing protein family. Each BTB-containing protein family forms a monophyletic group, identified with a colored background. K13 BTB is represented in red and clusters with the KCTD protein family of BTB-containing proteins. **c.** Patch of slowly evolving amino acid sites in a three-dimensional view of PfK13 BTB. The amino acid sites are labelled using the PfK13 sequence as reference. **d.** Superposition of the K13 BTB fold with two members of the KCTD protein family. The most conserved amino acid sites for each protein structure based on FuncPatch analysis are shown (magenta, orange and red for K13 BTB (white), SHKBP1 (green) and KCTD17 (cornflower blue), respectively). The patch of slowly evolving amino acid sites shared similarities between these three BTB-containing proteins, with three common positions: 357 on the B2-B3 loop and 397-398 on the B4-A4 loop.

To identify putative functional sites in K13 BTB, we examined whether a spatial correlation of the site-specific substitution rates *λ* is present in the K13 BTB-propeller tertiary structure. Despite a low standard deviation of substitution rates across amino acid sites, a significant spatial correlation was found with a log Bayes factor drastically > 8 and a 5 Å characteristic length (Table 2). The 10% most conserved sites predicted by FuncPatch formed one clearly bounded patch located at the surface of BTB (Fig. 3c). The BTB patch contained sites located at both B2-B3 and B4-A4 loops and at the A4 helix (positions/residues 355-358/NVGG, 397-399/DRD and 402-403/LF using the PfK13 sequence numbering; Fig. 3c). To test whether a similarly located, conserved patch was also found in the BTB domain of KCTD proteins, to which K13 BTB is the most similar, we inferred site-specific substitution rates *λ* from 124 and 139 orthologous sequences of SHKBP1 and KCTD17, respectively. For both proteins, the 10% most conserved BTB positions formed a patch that partially overlapped with the one of K13, with positions 357 (B2-B3 loop), 397 and 398 (B4-A4 loop) being shared between the three patches (PfK13 sequence numbering; Fig. 3d and Table 2). These positions are usually involved in BTB-BTB interactions in KCTDs^28^ or in BTB-Cullin interactions in some other BTB-containing protein families^27^, as in the X-ray structure of the Elongin C-Cullin2 complex (Supplementary Fig. 5). By functional annotation transfer, this indicates that these K13 BTB sites may be implied in some protein-protein interactions.

**Table 2.**
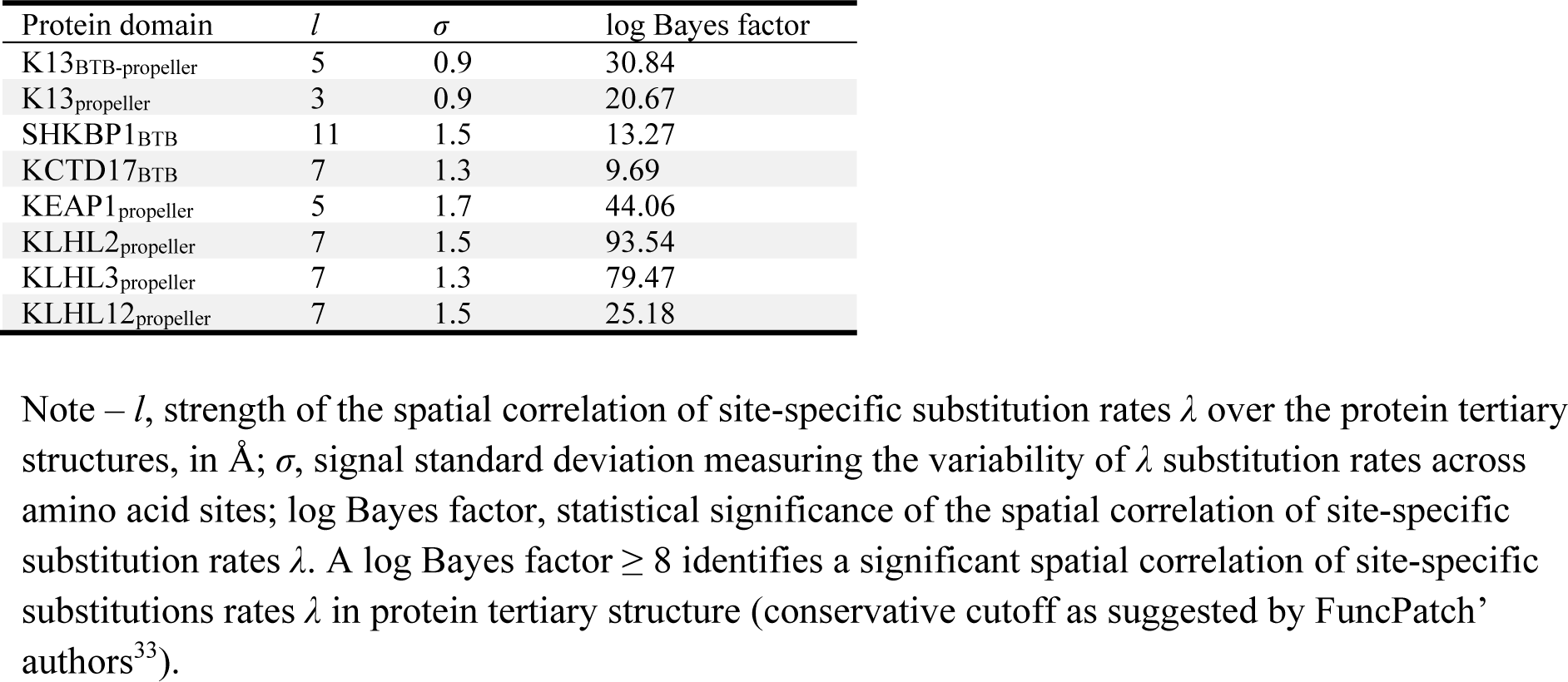
Spatial correlation of amino acid substitution rates in the BTB and propeller domains of K13 and other proteins estimated by FuncPatch analysis

| Protein domain | *l* | $\sigma$ | log Bayes factor |
| --- | --- | --- | --- |
| K13<sub>BTB</sub>-propeller | 5 | 0.9 | 30.84 |
| K13<sub>propeller</sub> | 3 | 0.9 | 20.67 |
| SHKBP1<sub>BTB</sub> | 11 | 1.5 | 13.27 |
| KCTD17<sub>BTB</sub> | 7 | 1.3 | 9.69 |
| KEAP1<sub>propeller</sub> | 5 | 1.7 | 44.06 |
| KLHL2<sub>propeller</sub> | 7 | 1.5 | 93.54 |
| KLHL3<sub>propeller</sub> | 7 | 1.3 | 79.47 |
| KLHL12<sub>propeller</sub> | 7 | 1.5 | 25.18 |

Note – *l*, strength of the spatial correlation of site-specific substitution rates $\lambda$ over the protein tertiary structures, in Å; $\sigma$ , signal standard deviation measuring the variability of $\lambda$ substitution rates across amino acid sites; log Bayes factor, statistical significance of the spatial correlation of site-specific substitution rates $\lambda$ . A log Bayes factor $\geq 8$ identifies a significant spatial correlation of site-specific substitutions rates $\lambda$ in protein tertiary structure (conservative cutoff as suggested by FuncPatch' authors<sup>33</sup>).

### The propeller domain of K13 exhibits a conserved shallow pocket

In Kelch-containing proteins, the propeller domain usually serves as the receptor for a substrate further ubiquitinated^23,24^. Before examining its conservation level, we first re-evaluated the architecture of the K13 propeller domain using its resolved 3D structure (PDB code 4YY8, chain A, unpublished). The PfK13 propeller structure is composed of six repeats of the Kelch motif (or blade)^8,22^. As expected, each blade is a β-sheet secondary structure involving four twisted antiparallel β-strands (numbered A to D). The innermost A strands line the central channel of the propeller fold whereas the outermost D strands are part of the propeller outside surface. The top face of the domain, containing the central channel opening, is formed by residues from several strands and AB and CD loops. The bottom face of propeller is composed of residues from the DA and BC loops and contains a shallow pocket, similar to other propeller structures^24^. Since there is no conventional definition for the shallow pocket delineation in the propeller fold, we defined it as the amino acids forming the surface plan of the pocket and protruding out of the plan (n = 19 positions; Fig. 4a and Supplementary Fig. 6).

**Fig 4.**
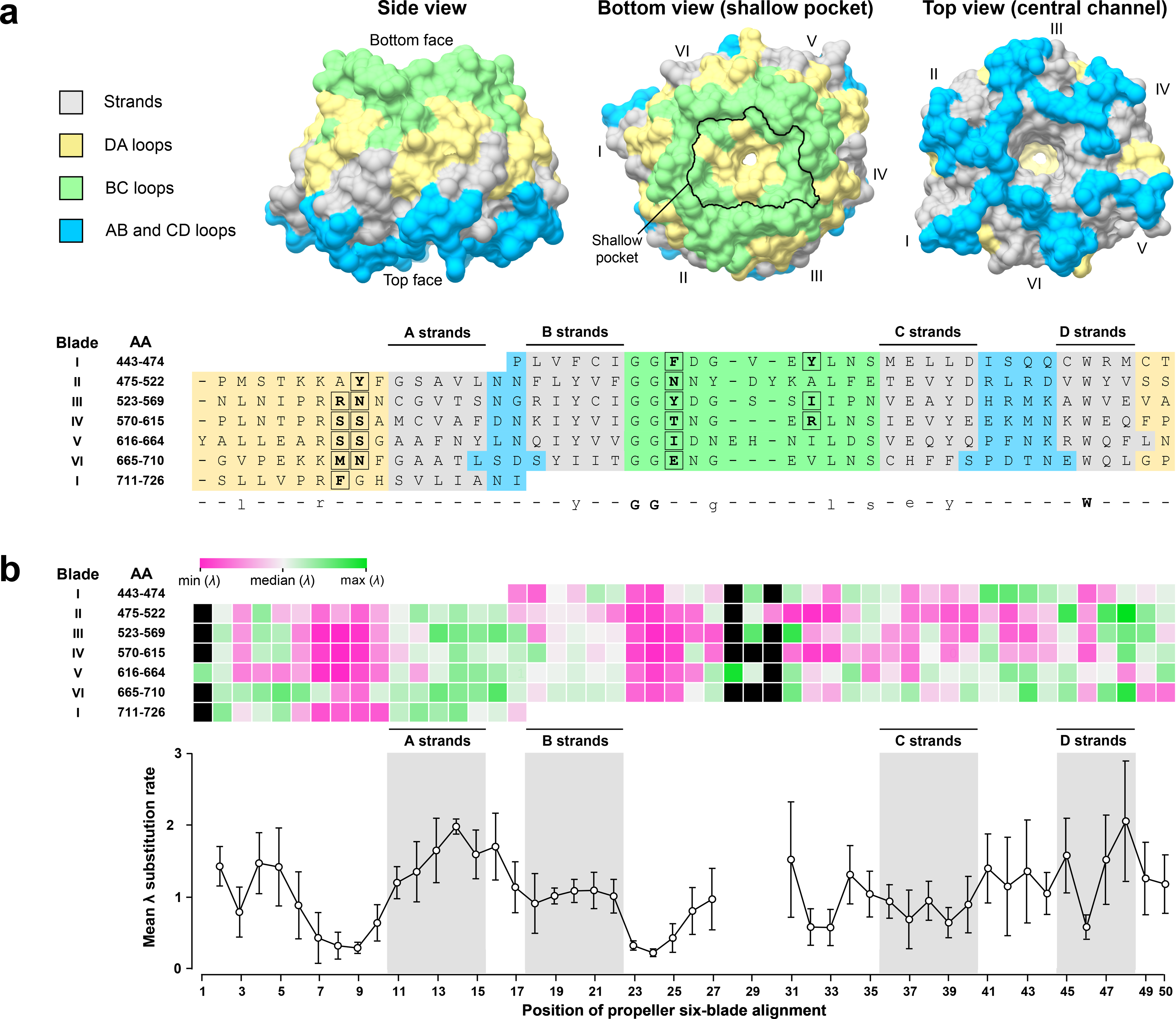
Structural organization of amino acid conservation across K13 propeller. **a.** Structure-based amino acid alignment of the six blade repeats of PfK13 propeller. The *x axis* shows the position in the structure-based amino acid alignment of the six blades; the left *y axis* shows the blade number, followed by the number of the first and last blade position. The strands of PfK13 propeller are colored in grey; the AB and CD loops forming the upper face of the propeller are colored in cyan; the BC and DA loops architecting the bottom face of the propeller are colored in yellow and green, respectively. A consensus sequence of the blades was defined: strict consensus amino acids are shown in bold capital letters, and highly conserved amino acids are shown in standard lowercase. A mapping of the different strands and loops onto the three-dimensional structure of PfK13 propeller is shown as surface above the structure-based amino acid alignment: side view (*left*), bottom view (*middle*) and upper view (*right*). The outline of the shallow pocket surface was delineated with a black line on the bottom view structure (*middle*) and the amino acid sites forming at the shallow pocket surface were surrounded and written in bold in the structure-based amino acid alignment. **b.** Mapping of the *λ* substitution rates onto the structure-based amino acid alignment of the six blade repeats of PfK13 propeller. *Upper graph*: heat map showing the *λ* substitution rate for each amino acid site. Black boxes correspond to the gaps in the structure-based amino acid alignment. The median *λ* substitution rate value was first calculated (white) to produce a scale ranging from the lowest (magenta) to the highest (green) site-specific substitution rate *λ*. *Lower graph*: Plot of the mean and 95% confidence interval of the *λ* substitution rates along the structure-based amino acid alignment of the six blades. The positions including one or more gaps were discarded.

To characterize the pattern of conservation within the domain, we superimposed the site-specific substitution rates *λ* inferred by FuncPatch onto an amino acid sequence alignment of the six blades (Fig. 4b), custom-produced from a structural alignment of the blades using PyMOL (Fig. 4a and Supplementary Fig. 7). We found that: *i)* the conservation level significantly differed between the six blades of propeller (*p* = 3.0 × 10^−3^, Kruskal-Wallis *H* test), with blade VI exhibiting the lowest conservation (Fig. 4b, Supplementary Fig. 8 and Supplementary Table 5); *ii)* loops were more conserved than strands (*p* = 5.0 × 10^−3^, Mann-Whitney *U* test; Fig. 4b and Supplementary Table 5); *iii)* the solvent-exposed A and D strands were less conserved than the buried B and C strands (*p* = 1.3 × 10^−6^, Kruskal-Wallis *H* test; Fig. 4b and Supplementary Table 5); and *iv)* the conservation level was the strongest at the blade positions 7-10 (DA loops) and 23-25 (BC loops; Fig. 4b), which altogether formed the surface and underlying layer of the shallow pocket in the PfK13 propeller tertiary structure (Fig. 4a). Of note, similar results were obtained with the *ω* estimates inferred under the best fitted PAML model (Supplementary Table 5).

With one exception (position PfK13 514), the 10% most conserved sites in K13 propeller (n = 29 out of 284) were all located at the bottom side of the propeller fold (Fig. 5a). Among them, the sites exposed at the surface (n = 9) formed a statistically significant patch located within the shallow pocket (Table 2 and Fig. 5a). The nineteen positions forming the shallow pocket of K13 propeller were significantly enriched in the 10% most conserved propeller amino acid sites (*p* = 1.6 × 10^−5^, chi-squared test; Fig. 5a, Tables 3 and 4). A similar trend was observed using the *ω* PAML values (Supplementary Fig. 9).

**Fig 5.**
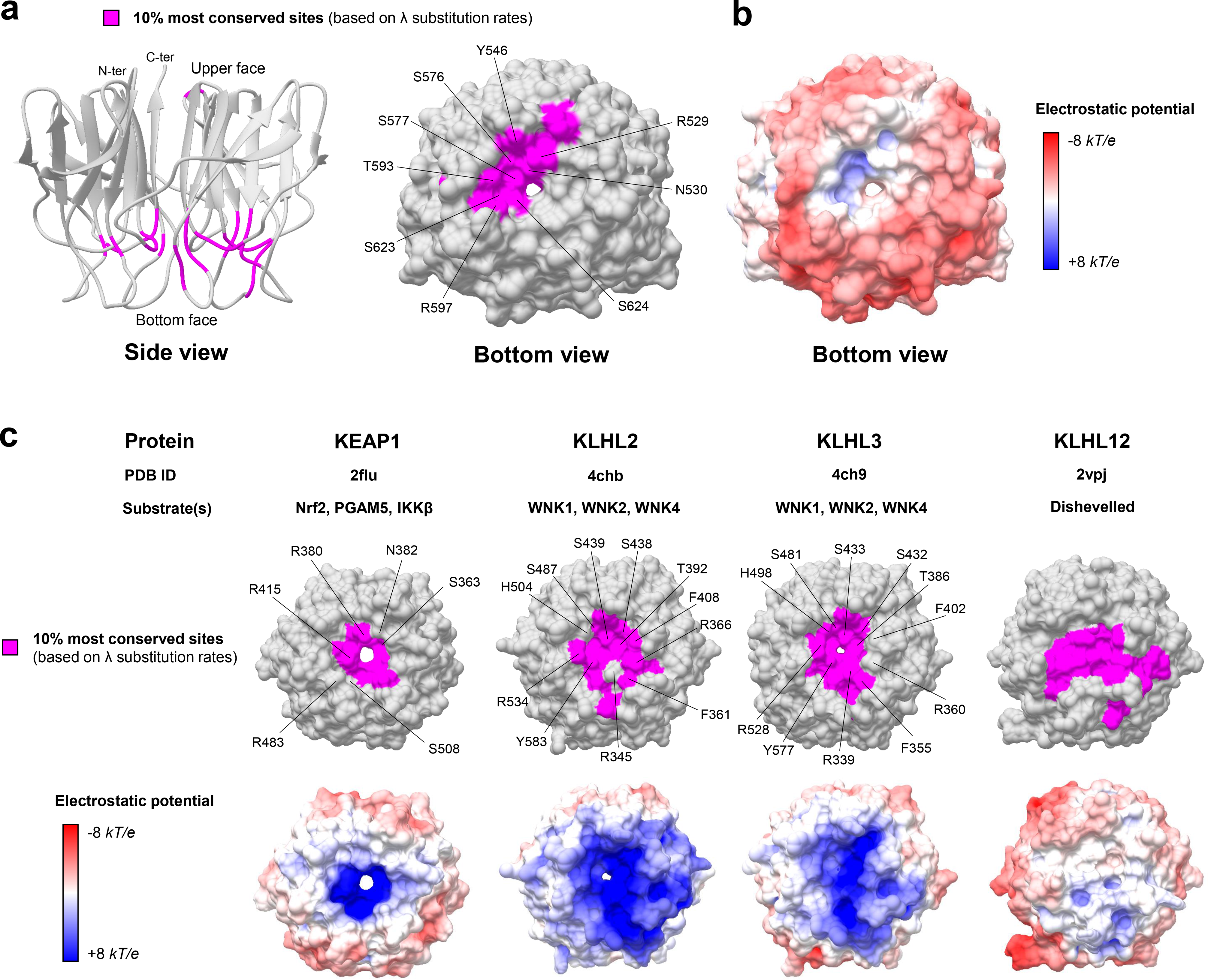
Conservation level and electrostatic potential across the propeller structures of K13 and other BTB-Kelch proteins. **a.** Location of the 10% most conserved amino acid sites (magenta) on the three-dimensional structure of PfK13 propeller. The conservation level of positions was defined using the site-specific substitution rates *λ* (Supplementary Data 1). The propeller structure is shown from the side view as cartoon (*left* structure) and from the bottom view as surface (*right* structure). The amino acid sites forming the surface of the shallow pocket and belonging to the 10% most conserved sites are labelled. **b.** Electrostatic surface potential of the PfK13 propeller structure, estimated with the APBS method. Electrostatic potential values are in units of *kT*/*e* at 298 K, on a scale of −8 *kT*/*e* (red) to +8 *kT*/*e* (blue). White color indicates a neutral potential. The missing charges were added using the *Add Charge* function implemented in USCF Chimera. **c.** Location of the 10% most conserved amino acid sites and electrostatic potential for KEAP1, KLHL2, KLHL3 and KLHL12 propeller structures. The color code and structure orientation are the same as for K13 in Fig. 4a, b. For KEAP1, KLHL2 and KLHL3, the key amino acids interacting with their respective protein substrates are labelled. The PDB IDs and protein substrates are provided above each propeller structure.

**Table 3.** Conservation level of the amino acids located at the surface of the K13 propeller shallow pocket

| Amino acid position | Reference amino acid (*P. fal*) | PfK13 mutant <sup>a</sup> | Sequence variation across species | | λ-based propeller rank n, (%) <sup>b</sup> |
| --- | --- | --- | --- | --- | --- |
|  |  |  | *Plasmodium* (n = 21) | Non-*Plasmodium* (n = 22) |  |
| 451 | F | — | F (21) | H (16); C (4); T (2); S(1); N (1) | 132 (46.48) |
| 456 | Y | — | Y (21) | H (12); I (7); P (2); Y (1) | 125 (44.01) |
| 482 | Y | S<sup>4</sup> (Africa) | Y (21) | Y (17); H (5) | 37 (13.03) |
| 498 | N | I<sup>80</sup> (Africa) | N (21) | Q (13); Y (7); H (2) | 44 (15.49) |
| **529** | **R** | **K<sup>80</sup> (Africa)** | **R (21)** | **R (22)** | **1 (0.35)** |
| **530** | **N** | **K/Y<sup>80</sup> (Africa)** | **N (21)** | **N (22)** | **9 (3.17)** |
| **546** | **Y** | — | **Y (21)** | **F (22)** | **13 (4.58)** |
| 551 | I | — | I (21) | I (19); V (3) | 86 (30.28) |
| **576** | **S** | **L<sup>81</sup> (Africa)** | **S (21)** | **S (22)** | **3 (1.06)** |
| **577** | **S** | **P<sup>4</sup> (Africa)** | **S (21)** | **S (22)** | **10 (3.52)** |
| **593** | **T** | **S<sup>10</sup> (Africa)** | **T (15); A (6)** | **T (22)** | **19 (6.69)** |
| **597** | **R** | — | **R (21)** | **R (22)** | **12 (4.23)** |
| **623** | **S** | **C<sup>8</sup> (Asia)** | **S (21)** | **S (22)** | **2 (0.70)** |
| **624** | **S** | — | **S (21)** | **A (15); G (7)** | **23 (8.10)** |
| 640 | I | V<sup>4</sup> (Africa) | I (21) | I (19); M (2); V(1) | 42 (14.79) |
| 671 | M | — | M (21) | M (17); L (3); I (1); F (1) | 73 (25.70) |
| 672 | N | — | N (21) | D (22) | 47 (16.55) |
| 688 | E | — | E (21) | Q (22) | 45 (15.85) |
| 717 | F | — | F (21) | Y (19); S (3) | 33 (11.62) |

Note – <sup>a</sup> None of the PfK13 mutant positions located at the surface of the K13 propeller shallow pocket was reported to confer the ART-R phenotype. All PfK13 mutants presented here have been described in one or two parasite samples from population surveys. <sup>b</sup> The rank attributed to the shallow pocket positions was based on the site-specific substitution rates $\lambda$ (Supplementary Data 1) estimated for the 284 studied propeller positions using FuncPatch<sup>33</sup>. The lower the rank, the more conserved was the position. Positions in bold are those belonging to the 10% most conserved positions of propeller.

**Table 4.** Distribution of the conserved propeller shallow pocket amino acids for K13 and some members of the BTB-Kelch protein family

| Protein | Conservation level <sup>a</sup> | Shallow pocket positions <sup>b</sup> | Others positions | *p* value |
| --- | --- | --- | --- | --- |
| PfK13<sub>propeller</sub> | Highly conserved<sup>10%</sup> | 9 | 20 | $1.60 \times 10^{-5}$ |
|  | Others | 10 | 245 |  |
| KEAP1<sub>propeller</sub> | Highly conserved<sup>10%</sup> | 12 | 17 | $1.44 \times 10^{-7}$ |
|  | Others | 12 | 244 |  |
| KLHL2<sub>propeller</sub> | Highly conserved<sup>10%</sup> | 13 | 16 | $1.26 \times 10^{-10}$ |
|  | Others | 6 | 251 |  |
| KLHL3<sub>propeller</sub> | Highly conserved<sup>10%</sup> | 12 | 17 | $1.36 \times 10^{-9}$ |
|  | Others | 6 | 251 |  |
| KLHL12<sub>propeller</sub> | Highly conserved<sup>10%</sup> | 14 | 15 | $2.94 \times 10^{-12}$ |
|  | Others | 5 | 255 |  |

Note – <sup>a</sup> The propeller positions were categorized into two groups: the *highly conserved* group contained the 10% most conserved positions according to the $\lambda$ substitution rates, the *other* group included the remaining propeller positions. <sup>b</sup> The shallow pocket of K13 propeller was defined as the positions forming the surface plan of the pocket and protruding out of the plan. A view and delineation of the PfK13 shallow pocket is provided in Supplementary Fig. 6. The same structural delineation of the propeller shallow pocket was applied to the four other BTB-Kelch propeller domains. Contingency tables were subjected to chi-squared test.

Using the PfK13 propeller structure as reference, we also identified two remarkable features of this conserved patch. First, it overlapped with a region of the shallow pocket harboring an electropositive surface potential energy, in contrast to the overall electronegative one of the propeller bottom surface (Fig. 5b). Second, it contained several arginine and serine residues strictly conserved in *Apicomplexa* (PfK13 R529, S576, S577, R597 and S623; Fig. 5a and Table 3), which are known to mediate protein-protein interactions in the pocket of other propeller domains^35,36^. Altogether, our evolutionary analysis of K13 propeller revealed that the shallow pocket is extremely conserved and may bind a substrate molecule.

### The conserved propeller patch predicted by FuncPatch is related to propeller binding activities in well-characterized BTB-Kelch proteins

To evaluate the reliability of FuncPatch to infer conserved functional sites in the context of propeller domains, we studied four other BTB-Kelch proteins found in mammals and which are functionally and structurally well-characterized (KEAP1, KLHL2, KLHL3 and KLHL12; Supplementary Fig. 10). All these proteins are known to bind a substrate protein through validated binding residues located in their propeller shallow pocket (substrates: Nrf2 for KEAP1^37^, WNK for KLHL2 and KLHL3^35^, and Dishevelled for KLHL12^38^). Using large sets of orthologous amino acid sequences (ranging from 129 sequences for KLHL12 to 162 sequences for KLHL2), a statistically significant spatial correlation of site-specific substitution rates *λ* was detected for each propeller fold (Table 2). In each case, the 10% most conserved propeller positions clustered in the shallow pocket (highest *p* = 1.4 × 10^−7^ for KEAP1, chi-squared test; Fig. 5c and Table 4). The shallow pocket of KEAP1, KLHL2 and KLHL3 propeller structures also showed a markedly electropositive surface potential energy while the one of KLHL12 was much more variable (Fig. 5c).

Therefore, the conserved functional sites predicted by FuncPatch in propeller domains from several BTB-Kelch proteins are consistent with the findings of previous experimental studies, demonstrating the reliability of our approach.

### Lack of association of ART-R mutations with some long-term evolutionary or structural facets of K13 propeller

Numerous non-synonymous mutations in PfK13 propeller segregating at variable frequencies have been reported to confer ART-R to *P. falciparum* parasites from SEA^4,8–11^. Here, we hypothesized that they may be associated with specific patterns of evolutionary-and/or structure-based parameters despite their large distribution across the propeller fold (Supplementary Fig 11). K13 propeller positions were classified as associated (n = 27) or not (n = 257) with an ART-R mutation, on the basis of the last World Health Organization (WHO) status report on ART-R^5^. No difference in the inter-species site-specific substitution rates *λ* was observed between the two groups (*p* = 0.96, Mann-Whitney *U* test; Supplementary Fig. 12). Importantly, we noted that no ART-R mutation has been reported at the positions located at the surface of the shallow pocket, although this trend was not statistically confirmed (*p* = 0.23, Fisher’s exact test; 0/19 ART-R mutations for the shallow pocket positions, and 27/265 ART-R mutations for the remaining K13 propeller positions). Two structure-based parameters associated with site evolutionary rate were also estimated: the relative solvent accessibility or RSA, and the side-chain weighted contact number or WCN_sc_^39^. Again, no association was found between these structural parameters and the propeller positions associated with ART-R mutations (*p* = 0.44 and *p* = 0.46, respectively, Mann-Whitney *U* test; Supplementary Fig. 12). In conclusion, none of the evolutionary- and structure-based parameters tested here were associated with ART-R K13 propeller positions.

## Discussion

Combining evolutionary and tertiary structure information provides a powerful and efficient way to gain insight into the functionality of protein sites^39^. The favored strategy usually screens sites that have evolved more rapidly than expected under a neutral model and interprets them as a signature of adaptive evolution corresponding to a gain of new function(s)^40,41^. Here, we focused on the most slowly evolving, 3D-bounded sites to identify highly conserved sub-regions of K13 across species evolution that are likely to play a functional role. Because of the extreme conservation of the *k13* gene^10^ and its essential function^14,15,42,43^, we made inter-rather than intra-species comparisons. In the context of a sustained and intense purifying selection operating in all annotated domains of K13, our analysis of sequence evolution coupled to BTB-propeller tertiary structure identified two patches of particularly slowly evolving sites.

The most striking conserved patch is found in the K13 propeller domain. It includes specific positions from the inter-blade DA and intra-blade BC loops, within a shallow pocket on the bottom face of the propeller structure. Several lines of evidence suggest that this patch contains functional sites. First, the shallow pocket is highly enriched in conserved positions, whereas solvent-exposed sites in proteins are usually less conserved than buried ones^39^. Second, the shallow pocket of well-characterized BTB-Kelch proteins directly mediates propeller’s binding activities (KEAP1, KLHL2, KLHL3, KLHL12)^24,35,36^ and are also predicted as conserved patches in our analyses. Finally, the conserved patches at the shallow pocket of BTB-Kelch proteins and K13 share interesting properties: they display a markedly electropositive surface potential energy and they are enriched in arginine and serine residues (PfK13 R529, R597, S576, S577 and S623, all strictly conserved in *Apicomplexa;* Table 3). In KEAP1, KLHL2 and KLHL3, the corresponding residues bind to acidic peptides derived from their substrates, Nrf2 and WNK, respectively^35,36^ (Fig. 5c). According to the electropositive region surrounding the K13 propeller conserved patch, the K13 substrate molecule(s) may harbor an acidic binding motif. Altogether, these results indicate that the shallow pocket of K13 propeller exhibits several properties of a binding surface, and we speculate that it may be critical for the recognition of substrate molecule(s).

In *P. falciparum*, PI3K is a likely candidate as it is immunoprecipitated with full-length PfK13, and its ubiquitination and proteasomal degradation are altered by the *pfk13* propeller C580Y mutation^17^. Another candidate may be the PK4 kinase which phosphorylates eIF2α, a key mediator of translation-mediated latency involved in ART-R^20^.

However, whether the *pfk13-mediated* ART-R mechanism in *P. falciparum* is related to the predicted binding activity of the propeller shallow pocket remains elusive. Based on the last WHO status report on ART-R^5^, none of the 27 validated or candidate ART-R mutations is located at the surface of the propeller shallow pocket. Several candidates are found in its underlayer (Supplementary Fig. 11) and some others have a preferential localization at positions proximal to the A and B strands^10^. Polymorphisms which have an uncharacterized phenotype were reported at 9 out of the 19 positions forming the shallow pocket, but found in only one or two parasite samples from population surveys (Table 3). Therefore, we speculate that amino acid changes at the positions contributing directly to the pocket surface are too damaging for the K13 native function to provide a long-term competitive advantage. In this context, rather than directly altering the binding residues, ART-R mutations may either induce long-range conformational changes propagating to the pocket surface or decrease the propeller domain stability, as reported for pathogenic mutations in KEAP1 W544C and KLHL3 S410L, respectively^44,45^. Furthermore, putative propeller conformational changes associated with ART-R mutations might be induced by specific cellular conditions, for example increased oxidative stress^46^.

BTB appeared here as the most conserved domain of K13 during *Apicomplexa* evolution, and therefore likely carries critical activities. It most resembles the BTB of the KCTD protein family in primary sequence, tertiary structure and short domain size. The shortened BTB of KCTDs could still mediate protein oligomerization^28,47^, consistent with the dimer observed in the solved PfK13 BTB-propeller crystal structures (PDB codes 4YY8 and 4ZGC, unpublished). K13 BTB harbors a predicted, functional patch – located at the B2-B3 and B4-A4 loops and at the A4 helix – that overlaps with the one of KCTD proteins, suggesting that they share similar functional sites. In KCTDs, these sites make BTB-BTB contacts in tetrameric or pentameric assemblies when BTB is solved as an isolated domain^28^. However, the PfK13 BTB-propeller structure forms a dimer and none of the highly conserved positions make BTB-BTB contacts (PDB codes 4YY8 and 4ZGC, unpublished). Finally, amino acids of the B4-A4 loop (corresponding to positions 397-399 in PfK13) are exposed at the BTB-Cullin binding interface in several solved complexes^27^. These discrepancies in the role of the predicted BTB patch could be due to the fact that additional domains (such as propeller) or partner proteins might constraint the folding of the BTB domain into oligomers or complexes. Altogether, data from the literature however support that the BTB predicted patch of K13 mediates protein-protein interactions, possibly with a Cullin protein.

Interestingly, K13 also contains a highly conserved CCC domain, located before BTB, which therefore likely carries critical activities. Consistent with this hypothesis, three *pfk13* mutations conferring a moderate ART-R are located in CCC (PfK13 R239Q, E252Q and D281V)^3,11,48^. Coiled-coils are ubiquitous protein-protein interaction domains composed of two or more α-helices coiling together^49^. A CCC domain was reported in a few Kelch-containing proteins (including some KCTDs) involved in cell morphogenesis but these CCC have a different domain organization than the one of K13^22^. The CCC of K13 may participate in K13 oligomerization and/or serve as a binding interface with other proteins^50^.

In conclusion, through evolutionary and structural analyses, we identified the shallow pocket of the K13 propeller domain as a candidate surface for binding a substrate molecule. We also detected in the BTB domain of K13 a conserved patch of sites that are involved in protein-protein interactions in known BTB-Cullin and BTB-BTB complexes. Efforts should now focus on the identification of molecule(s) that bind(s) to the pocket of K13 propeller, which may help clarify the link between K13 function and ART-R.

## Materials and Methods

### Collection of *k13* orthologous sequences from genomic databases

The amino acid sequence of PfK13 (PlasmoDB code PF3D7_1343700) was queried against the specialized eukaryotic pathogen database (EuPathDB release 33)^51^ and the NCBI non-redundant protein database using blastp and tblastn searches^52^ (BLOSUM62 matrix). A protein was considered as a likely orthologous sequence if the sequence identity was ≥ 30% and the e-value below the 10^−3^ cutoff. Forty three K13 sequences – and corresponding *k13* cDNA sequences – were retrieved from distinct *Apicomplexa* species including 21 *Plasmodium* species. A detailed bioinformatics analysis was performed on each protein sequence to confirm the presence of the three annotated domains of K13 (CCC, BTB and propeller) using InterPro^53^.

### *k13* sequence alignment

Considering the greater divergence of coding nucleotide sequences as compared to protein sequences due to the genetic code redundancy, a K13 protein sequence alignment was first generated using Mafft version 7^54^ (E-INS-I strategy with BLOSUM62 scoring matrix, gap opening penalty 2.0 and offset 0.1). The output alignment was visually inspected and manually edited with BioEdit v7.2.5^55^. The positions containing gaps in at least 30% of all sequences were removed, as suggested by PAML’ authors^31^. Then, the *k13* nucleotide sequence alignment was generated with PAL2NAL^56^ using the cleaned K13 amino acid alignment as template.

### Phylogenetic analysis of *k13*

The phylogenetic relationships of *k13* nucleotide sequences were inferred using the maximum-likelihood method implemented in PhyML v3.0^57^, after determining the best-fitting nucleotide substitution model using the Smart Model Selection (SMS) package^58^. A general time-reversible model with optimized equilibrium frequencies, gamma distributed among-site rate variation and estimated proportion of invariable sites (GTR + *G* + *I*) was used, as selected by the Akaike Information Criterion. The nearest neighbor interchange approach was chosen for tree improving, and branch supports were estimated using the approximate likelihood ratio aLRT SH-like method^59^. The *k13* phylogeny was rooted using *Cryptosporidia* species as outgroup.

### Molecular evolutionary analysis of *k13*

To investigate the evolutionary regime that has shaped the *k13* protein-coding DNA sequence during species evolution, we analyzed the non-synonymous (*d*_N_) to synonymous *d*_S_) substitution rate ratio *ω* (= *d*_N_/*d*_S_), estimated by maximum-likelihood using the codeml tool from PAML v.4.8^31,61^. *ω* provides a sensitive measure of selective pressure at the amino acid level by comparing substitution rates with statistical distribution and considering the phylogenetic tree topology. Typically, *ω* < 1 indicates purifying selection, while *ω* = 1 and *ω* > 1 indicate neutral evolution and positive selection, respectively.

The heterogeneity of *ω* among lineages of the *k13* phylogenetic tree (branch models) was tested by comparing the free-ratio (FR) model, which assumes as many *ω* parameters as the number of branches in the tree, to the one-ratio (M0) model which supposes only one *ω* value for all branches^62,63^.

The variation of *ω* among codon sites was then evaluated using codon models M1a, M2a, M3, M7 and M8^62,63^. M1a allows codon sites to fall into two site classes, either with *ω* < 1 (purifying selection) or *ω* = 1 (neutral evolution), whereas model M2a extends model M1a with a further site class as *ω* > 1 (positive selection). Model M3 includes a discrete distribution of independent *ω* with *k* classes of sites (*k* = [3, 4, 5] in this study), with *ω* values and corresponding proportions estimated from the dataset. Model M7 assumed a *β*-distribution of ten *ω* ratios limited to the interval [0, 1] with two shape parameters *p* and *q*, whereas model M8 adds an additional site class with *ω* possibly > 1 as M2a does. The heterogeneity of *ω* across codon sites was tested by comparison of models M0-M3, while comparison of paired models M1a-M2a and M7-M8 allowed to detect positive selection^62^.

Model comparisons were made using likelihood ratio tests (LRTs)^64^. For each of the LRTs, twice the log-likelihood difference between alternative and null models (2ΔlnL) was compared to critical values from a chi-squared distribution with degrees of freedom equal to the difference in the number of estimated parameters between both models^40^. Candidate sites for positive selection were pinpointed using the Bayes empirical Bayes (BEB) inference which calculates the posterior probability that each codon site falls into a site class affected by positive selection (in models M2a and M8), as described by Yang and colleagues^65^. For model M3, in which no BEB approach is implemented yet, the naive empirical Bayes (NEB) approach was used to identify those sites evolving under positive selection.

Three codon substitution models were used and compared for all models: F1×4 and F3×4, which assume equal nucleotide frequencies and individual codon frequencies at all codon positions, respectively, and the parameter-rich model F61, which estimates codon frequencies separately for each codon^61,66^. Since the three codon substitution models yielded similar results (Supplementary Fig. 13), we only presented those obtained with the most widely used F3×4 codon model. The analyses were run multiple times with different *ω* starting values to check the consistency of the results.

For PAML model M3 with *k* site classes of *ω* ratios, the posterior mean of *ω* value at each codon site was calculated as the average of the *ω* ratios across the *k ω* site classes weighted by their posterior probabilities^31^.

In addition to *k13*, four other Kelch protein-coding sequences were considered to compare their *ω* with those estimated for the whole *Plasmodium* proteome. We used the *ω* values previously estimated by Jeffares and colleagues^32^ with PAML under the one-ratio model for each of the 3,256 orthologous protein-coding genes among six *Plasmodium* species: *P. falciparum*, *P. berghei*, *P. chabaudi*, *P. vivax*, *P. yoelii* and *P. knowlesi*. A full description of the procedure is presented in the original paper^32^.

### Inferring site-specific substitution rates considering their spatial correlation in the K13 BTB-propeller tertiary structure

Most methods – including PAML – assume that site-specific substitution rates are independently distributed across sites^67^. However, it is widely acknowledged that amino acids located close to each other in protein tertiary structures are more likely to carry out similar functions, suggesting a site interdependence in amino acid sequence evolution attributed to tertiary structure^67,68^. Consequently, the substitution rate at the protein level (named *λ* in this study) was inferred using the FuncPatch server^33^. FuncPatch requires an amino acid sequence alignment, a phylogenetic tree and a protein tertiary structure to estimate the conservation level during species evolution and the characteristic scale (in Å) of spatially correlated site-specific substitution rates *λ*. We used the X-ray structure at 1.81 Å resolution of PfK13 BTB-propeller as the reference structure which does not contain the conserved CCC domain (PDB code 4YY8, unpublished). Beforehand, a Ramachandran analysis was performed to validate the quality of the structure using MolProbity^69^: 96.9% and 3.1% of the amino acids were in favored and allowed regions, respectively, and there were no outliers. FuncPatch only accepts monomeric proteins as input whereas BTB-propeller of PfK13 dimerizes in crystal structure. To take into account the dimeric organization of PfK13, its tertiary structure was edited using customized python scripts (Python v2.7.13) in order to merge the two monomers (chains A and B) and the K13 sequence was duplicated in the K13 protein sequence alignment. The analysis was also done using either one of the other monomeric BTB-propeller tertiary structure and also using a disulfide-bonded version of PfK13 BTB-propeller (PDB code 4ZGC, unpublished). All these control analyses yielded similar results (data not shown). The spatial correlation of the site-specific substitution rates *λ* in the K13 tertiary structure was tested using a Bayesian model comparison, where a null model (model 0), in which no spatial correlation of site-specific substitution rates *λ* is present, is compared to the alternative model (model 1). As suggested by FuncPatch’ authors, the spatial correlation was considered as significant if the estimated log Bayes factor (model 1 versus model 0) was larger than 8 in the dataset (conservative cutoff)^33^.

### Delineation of K13 propeller blades and secondary structures

The propeller domain of PfK13 is composed of six blades having slightly different amino acid lengths. To get an accurate blade alignment at the primary amino acid sequence level, we first sought to align the six blade structures. The PDB propeller structure was obtained from the PfK13 BTB-propeller structure (PDB code 4YY8, chain A, unpublished) and was then divided into six parts, each one containing the atomic coordinates of one blade. The six blade structures were then aligned by minimizing the RMSD of atomic positions using the *align* function in PyMOL Molecular Graphics System^70^ so as to identify the amino acids from the six blades that are located at exactly the same blade position. This structure alignment was then used to align the six blades at the primary amino acid sequence level. The delineation of the strands and loops was obtained directly from the PDB file (PDB code 4YY8, chain A, unpublished).

### Definition of ART-R mutations

We used the last status report on ART-R provided by the WHO to classify the positions of the PfK13 propeller domain as associated or not with an ART-R mutation^5^.

### Evolutionary analysis of the BTB and propeller domains in other BTB-and Kelch-containing proteins

To characterize the BTB domain of K13, we arbitrarily retrieved some members belonging to the main BTB-containing protein families (BTB-ZF, BTB-Kelch, RhoBTB, BTB-NPH3, MATH-BTB, KCTD, KCNA and SKP1 and Elongin C proteins; full list provided in Supplementary Table 6). A multiple protein alignment was generated using Mafft version 7^54^ (default parameters) and was then manually edited with BioEdit v7.2.5^55^ to retain only the region referring to the BTB core fold. The phylogenetic relationships were inferred with the PhyML^57^ procedure using the best-fitting protein substitution model as determined by the SMS package^58^.

For further comparisons with the K13 BTB and propeller domains, site-specific substitution rates *λ* were inferred with FuncPatch for the BTB and propeller domains of several mammalian KCTD and BTB-Kelch proteins, respectively. In the present study, the proteins were selected on the basis of their sequence homology with K13, the availability of a solved 3D structure, and their known implication in a Cullin-RING E3 ligase complex as suspected for K13. In addition, only well-characterized ligand-binding function and the presence of a six-bladed propeller structure similar to the one of K13 were considered for BTB-Kelch proteins. After a careful review of the literature, we selected two KCTD proteins: SHKBP1 (UniProt code Q8TBC3) which regulates the epidermal growth factor receptor (EGFR) signaling pathway^71^; and KCTD17 (Q8N5Z5) which mediates the ubiquitination and proteasomal degradation of the ciliogenesis down-regulation TCHP protein^72^. Considering BTB-Kelch proteins, we focused on: KEAP1 (Q14145) which interacts with its client protein Nrf2 for the induction of cytoprotective responses to oxidative stress^37^; KLHL2 (O95198) and KLHL3 (Q9UH77) which both participate in the ubiquitination and degradation of WNK substrates regulating blood pressure^35^; and KLHL12 (Q53G59) which negatively regulates the WNT-beta-catenin pathway through the degradation of Dishevelled proteins^38^. First, each *Homo sapiens* KCTD and BTB-Kelch sequence was successively submitted as query sequence for a blastp search^52^ (BLOSUM62 scoring matrix, max target sequences fixed at 1,000) against the NCBI non-redundant protein database to retrieve orthologous sequences from a large amount of species. The output lists were then filtered according to specific criteria so as to keep only sequences having an unambiguous description (*i.e.* a description that includes the name of the queried KCTD or BTB-Kelch protein), and that aligned with ≥ 80% sequence coverage and had ≥ 60% sequence identity with the query sequence. The multiple protein alignment of each set of orthologous sequences was then generated using Mafft version 7^54^ (E-INS-I strategy with BLOSUM62 scoring matrix, gap opening penalty 2.0 and offset 0.1). A second filtering step was performed to remove incomplete or miss-annotated sequences, *i.e.* the sequences that did not contain all the annotated domains (using the domain annotation automatically generated by the Uniprot Knowledgebase) and/or that included a gapped position located in one of the annotated domains. The final multiple protein alignments included: *i*) 124 sequences × 103 aligned positions for SHKBP1; *ii*) 139 sequences × 102 aligned positions for KCTD17; *iii*) 135 sequences × 285 aligned positions for KEAP1; *iv*) 162 sequences × 286 aligned positions for KLHL2; *v*) 158 sequences × 286 aligned positions for KLHL3; and *vi*) 129 sequences × 289 aligned positions for KLHL12. The full list of orthologous sequences used for each mammalian KCTD and BTB-Kelch protein is provided in Supplementary Table 7. Then, the phylogenetic relationships were inferred using PhyML^57^ after determining the best-fitting protein substitution model with the SMS package^58^. The 3D structures of KCTD BTB and BTB-Kelch propeller domains were retrieved from the PDBsum database under the following accession numbers: 4CRH for SHKBP1 (resolution: 1.72 Å)^28^, 5A6R for KCTD17 (resolution: 2.85 Å)^28^, 2FLU for KEAP1 (resolution: 1.50 Å, in complex with a Nrf2 peptide)^73^, 4CHB for KLHL2 (resolution: 1.56 Å, in complex with a WNK4 peptide)^35^, 4CH9 for KLHL3 (resolution: 1. 84 Å, in complex with a WNK4 peptide)^35^, and 2VPJ for KLHL12 (resolution: 1.85 Å)^24^. Beforehand, the quality of each structure was validated using MolProbity^69^: none of the structures had amino acids identified as outliers, and approximately 98% of the amino acids of each structure were in favored regions.

### Evaluation of structural properties

The electrostatic potential energy of each propeller structure was calculated using the Adaptive Poisson-Boltzmann Solver (APBS) method^74^. Beforehand, the required pqr input files were prepared using PDB2PQR v.2.1.1^75^. The missing charges were added using the *Add Charge* function implemented in USCF Chimera^76^. A grid-based method was used to solve the linearized Poisson-Boltzmann equation at 298 K, with solute (protein) and solvent dielectric constant values fixed at 2 and 78.5, respectively. The contact surface selection was mapped using a radius of 1.4 Å in a scale of −8 *kT/e* to +8 *kT/e.*

The relative solvent accessibility (RSA) was estimated as the accessible surface area of amino acids using DSSP^77^, then normalized with the maximal accessible surface area of each amino acid^78^. The side-chain weighted contact number (WCN_sc_) of each amino acid was calculated using a customized python script provided by Sydykova and colleagues^79^. All structural properties were assessed using the aforementioned PDB files.

### Structure visualization

All molecular drawings were generated using the UCSF Chimera software^76^.

### Statistical analyses

Substitution rates among partitions were compared using non-parametric Mann-Whitney *U* or Kruskal-Wallis *H* tests. When focusing on the propeller shallow pocket, contingency tables were produced and statistically tested with chi-squared test. We used *p* < 0.05 as the cutoff for significance in all statistical tests.

## Acknowledgements

We thank YF. Huang and GB. Golding for help with FuncPatch analysis. We thank O. Mercereau-Puijalon, M. Miteva, and R. Duval for helpful discussions. We thank A. Sissoko and F. Palstra for proofreading the manuscript. R. Coppée is supported by PhD grant funding from the École Doctorale MTCI.

## Author contributions

R.C., A.S. and J.C. designed the study. D.J. provided data. R.C. performed most of the analyses. R.C., A.S., and J.C. contributed to interpretations. All authors have participated in paper writing and approved it prior to submission.

## Competing interests

The authors declare no competing financial interests.

## References

1. Dondorp, A. M. et al. Artemisinin resistance in Plasmodium falciparum malaria. N. Engl. J. Med. 361, 455–467 (2009).

2. Duru, V. et al. Plasmodium falciparum dihydroartemisinin-piperaquine failures in Cambodia are associated with mutant K13 parasites presenting high survival rates in novel piperaquine in vitro assays: retrospective and prospective investigations. BMC Med. 13, 305 (2015).

3. Phyo, A. P. et al. Declining efficacy of artemisinin combination therapy against P. falciparum malaria on the Thai-Myanmar border (2003-2013): the role of parasite genetic factors. Clin. Infect. Dis. 63, 784–791 (2016).

4. Ménard, D. et al. A worldwide map of Plasmodium falciparum K13-propeller polymorphisms. N. Engl. J. Med. 374, 2453–2464 (2016).

5. WHO. Status report on artemisinin and ACT resistance, april 2017. http://www.who.int/malaria/publications/atoz/artemisinin-resistance-april2017/en/ (2017).

6. Lu, F. et al. Emergence of indigenous artemisinin-resistant Plasmodium falciparum in Africa. N. Engl. J. Med. 376, 991–993 (2017).

7. Ikeda, M. et al. Artemisinin-resistant Plasmodium falciparum with high survival rates, Uganda, 2014-2016. Emerging Infect. Dis. 24, 718–726 (2018).

8. Ariey, F. et al. A molecular marker of artemisinin-resistant Plasmodium falciparum malaria. Nature 505, 50–55 (2014).

9. Straimer, J. et al. K13-propeller mutations confer artemisinin resistance in Plasmodium falciparum clinical isolates. Science 347, 428–431 (2015).

10. MalariaGEN Plasmodium falciparum Community Project. Genomic epidemiology of artemisinin resistant malaria. Elife 5, pii: e08714 (2016).

11. Anderson, T. J. C. et al. Population parameters underlying an ongoing soft sweep in Southeast Asian malaria parasites. Mol. Biol. Evol. 34, 131–144 (2017).

12. Cerqueira, G. C. et al. Longitudinal genomic surveillance of Plasmodium falciparum malaria parasites reveals complex genomic architecture of emerging artemisinin resistance. Genome Biol. 18, 78 (2017).

13. Witkowski, B. et al. Novel phenotypic assays for the detection of artemisinin-resistant Plasmodium falciparum malaria in Cambodia: in-vitro and ex-vivo drug-response studies. Lancet Infect. Dis. 13, 1043–1049 (2013).

14. Birnbaum, J. et al. A genetic system to study Plasmodium falciparum protein function. Nat. Methods 14, 450–456 (2017).

15. Zhang, M. et al. Uncovering the essential genes of the human malaria parasite Plasmodium falciparum by saturation mutagenesis. Science 360, pii: eaap7847 (2018).

16. The Gene Ontology Consortium. Expansion of the gene ontology knowledgebase and resources. Nucleic Acids Res. 45, D331–D338 (2017).

17. Mbengue, A. et al. A molecular mechanism of artemisinin resistance in Plasmodium falciparum malaria. Nature 520, 683–687 (2015).

18. Dogovski, C. et al. Targeting the cell stress response of Plasmodium falciparum to overcome artemisinin Resistance. PLoS Biol. 13, e1002132 (2015).

19. Mok, S. et al. Population transcriptomics of human malaria parasites reveals the mechanism of artemisinin resistance. Science 347, 431–435 (2015).

20. Zhang, M. et al. Inhibiting the Plasmodium eIF2α kinase PK4 prevents artemisinin-induced latency. Cell Host & Microbe 22, 766–776.e4 (2017).

21. Bhattacharjee, S. et al. Remodeling of the malaria parasite and host human red cell by vesicle amplification that induces artemisinin resistance. Blood 131, 1234–1247 (2018).

22. Adams, J., Kelso, R. & Cooley, L. The kelch repeat superfamily of proteins: propellers of cell function. Trends Cell Biol. 10, 17–24 (2000).

23. Dhanoa, B. S., Cogliati, T., Satish, A. G., Bruford, E. A. & Friedman, J. S. Update on the Kelch-like (KLHL) gene family. Hum. Genomics 7, 13 (2013).

24. Canning, P. et al. Structural basis for Cul3 protein assembly with the BTB-Kelch family of E3 ubiquitin ligases. J. Biol. Chem. 288, 7803–7814 (2013).

25. Tilley, L., Straimer, J., Gnädig, N. F., Ralph, S. A. & Fidock, D. A. Artemisinin action and resistance in Plasmodium falciparum. Trends Parasitol. 32, 682–696 (2016).

26. Geyer, R., Wee, S., Anderson, S., Yates, J. & Wolf, D. A. BTB/POZ domain proteins are putative substrate adaptors for Cullin 3 ubiquitin ligases. Mol. Cell 12, 783–790 (2003).

27. Stogios, P. J., Downs, G. S., Jauhal, J. J., Nandra, S. K. & Privé, G. G. Sequence and structural analysis of BTB domain proteins. Genome Biol. 6, R82 (2005).

28. Pinkas, D. M. et al. Structural complexity in the KCTD family of Cullin3-dependent E3 ubiquitin ligases. Biochem. J. 474, 3747–3761 (2017).

29. Furukawa, M. & Xiong, Y. BTB protein Keap1 targets antioxidant transcription factor Nrf2 for ubiquitination by the Cullin 3-Roc1 ligase. Mol. Cell. Biol. 25, 162–171 (2005).

30. Shibata, S., Zhang, J., Puthumana, J., Stone, K. L. & Lifton, R. P. Kelch-like 3 and Cullin 3 regulate electrolyte homeostasis via ubiquitination and degradation of WNK4. Proc. Natl. Acad.Sci. U.S.A. 110, 7838–7843 (2013).

31. Yang, Z. PAML 4: phylogenetic analysis by maximum likelihood. Mol. Biol. Evol. 24, 1586–1591 (2007).

32. Jeffares, D. C., Tomiczek, B., Sojo, V. & dos Reis, M. A beginners guide to estimating the non-synonymous to synonymous rate ratio of all protein-coding genes in a genome. Methods Mol. Biol. 1201, 65–90 (2015).

33. Huang, Y.-F. & Golding, G. B. FuncPatch: a web server for the fast Bayesian inference of conserved functional patches in protein 3D structures. Bioinformatics 31, 523–531 (2015).

34. Huang, Y.-F. & Golding, G. B. Phylogenetic Gaussian process model for the inference of functionally important regions in protein tertiary structures. PLoS Comput. Biol. 10, e1003429 (2014).

35. Schumacher, F. R., Sorrell, F. J., Alessi, D. R., Bullock, A. N. & Kurz, T. Structural and biochemical characterization of the KLHL3-WNK kinase interaction important in blood pressure regulation. Biochem. J. 460, 237–246 (2014).

36. Canning, P., Sorrell, F. J. & Bullock, A. N. Structural basis of Keap1 interactions with Nrf2. Free Radic. Biol. Med. 88, 101–107 (2015).

37. Cullinan, S. B. et al. Nrf2 Is a Direct PERK substrate and effector of PERK-dependent cell survival. Mol. Cell. Biol. 23, 7198–7209 (2003).

38. Angers, S. et al. The KLHL12-Cullin-3 ubiquitin ligase negatively regulates the Wnt-β-catenin pathway by targeting Dishevelled for degradation. Nat. Cell Biol. 8, 348–357 (2006).

39. Echave, J., Spielman, S. J. & Wilke, C. O. Causes of evolutionary rate variation among protein sites. Nat. Rev. Genet. 17, 109–121 (2016).

40. Yang, Z., Nielsen, R., Goldman, N. & Pedersen, A. M. Codon-substitution models for heterogeneous selection pressure at amino acid sites. Genetics 155, 431–449 (2000).

41. Sawyer, S. L., Wu, L. I., Emerman, M. & Malik, H. S. Positive selection of primate TRIM5alpha identifies a critical species-specific retroviral restriction domain. Proc. Natl. Acad. Sci. U.S.A. 102, 2832–2837 (2005).

42. Sidik, S. M. et al. A genome-wide CRISPR screen in Toxoplasma identifies essential apicomplexan genes. Cell 166, 1423–1435.e12 (2016).

43. Bushell, E. et al. Functional profiling of a plasmodium genome reveals an abundance of essential genes. Cell 170, 260–272.e8 (2017).

44. Hast, B. E. et al. Cancer-derived mutations in KEAP1 impair NRF2 degradation but not ubiquitination. Cancer Res. 74, 808–817 (2014).

45. Mori, Y. et al. Decrease of WNK4 ubiquitination by disease-causing mutations of KLHL3 through different molecular mechanisms. Biochem. Biophys. Res. Commun. 439, 30–34 (2013).

46. Haldar, K., Bhattacharjee, S. & Safeukui, I. Drug resistance in Plasmodium. Nat. Rev. Microbiol. 16, 156–170 (2018).

47. Correale, S. et al. Molecular organization of the cullin E3 ligase adaptor KCTD11. Biochimie 93, 715–724 (2011).

48. Boullé, M. et al. Artemisinin-resistant Plasmodium falciparum K13 mutant alleles, Thailand-Myanmar border. Emerg. Infect. Dis. 22, 1503–1505 (2016).

49. Hartmann, M. D. Functional and structural roles of coiled coils. Subcell. Biochem. 82, 63–93 (2017).

50. de Moura, T. R. et al. Prp19/Pso4 is an autoinhibited ubiquitin ligase activated by stepwise assembly of three splicing factors. Mol. Cell 69, 979–992.e6 (2018).

51. Aurrecoechea, C. et al. EuPathDB: the eukaryotic pathogen genomics database resource. Nucleic Acids Res. 45, D581–D591 (2017).

52. Altschul, S. F., Gish, W., Miller, W., Myers, E. W. & Lipman, D. J. Basic local alignment search tool. J. Mol. Biol. 215, 403–410 (1990).

53. Hunter, S. et al. InterPro: the integrative protein signature database. Nucleic Acids Res. 37, D211–D215 (2009).

54. Katoh, K. & Standley, D. M. MAFFT multiple sequence alignment software version 7: improvements in performance and usability. Mol. Biol. Evol. 30, 772–780 (2013).

55. Hall, T. BioEdit: a user-friendly biological sequence alignment editor and analysis program for Windows 95/98/NT. Nucleic Acids Symp. Ser. 41, 95–98 (1999).

56. Suyama, M., Torrents, D. & Bork, P. PAL2NAL: robust conversion of protein sequence alignments into the corresponding codon alignments. Nucleic Acids Res. 34, W609–W612 (2006).

57. Guindon, S. et al. New algorithms and methods to estimate maximum-likelihood phylogenies: assessing the performance of PhyML 3.0. Syst. Biol. 59, 307–321 (2010).

58. Lefort, V., Longueville, J.-E. & Gascuel, O. SMS: smart model selection in PhyML. Mol. Biol. Evol. 34, 2422–2424 (2017).

59. Anisimova, M., Bielawski, J. P. & Yang, Z. Accuracy and power of the likelihood ratio test in detecting adaptive molecular evolution. Mol. Biol. Evol. 18, 1585–1592 (2001).

60. Parfrey, L. W., Lahr, D. J. G., Knoll, A. H. & Katz, L. A. Estimating the timing of early eukaryotic diversification with multigene molecular clocks. Proc. Natl. Acad. Sci. U.S.A. 108, 13624–13629 (2011).

61. Goldman, N. & Yang, Z. A codon-based model of nucleotide substitution for protein-coding DNA sequences. Mol. Biol. Evol. 11, 725–736 (1994).

62. Nielsen, R. & Yang, Z. Likelihood models for detecting positively selected amino acid sites and applications to the HIV-1 envelope gene. Genetics 148, 929–936 (1998).

63. Yang, Z. Likelihood ratio tests for detecting positive selection and application to primate lysozyme evolution. Mol. Biol. Evol. 15, 568–573 (1998).

64. Vuong, Q. H. Likelihood ratio tests for model selection and non-nested hypotheses. Econometrica 57, 307–333 (1989).

65. Yang, Z., Wong, W. S. W. & Nielsen, R. Bayes empirical bayes inference of amino acid sites under positive selection. Mol. Biol. Evol. 22, 1107–1118 (2005).

66. Muse, S. V. & Gaut, B. S. A likelihood approach for comparing synonymous and nonsynonymous nucleotide substitution rates, with application to the chloroplast genome. Mol. Biol. Evol. 11, 715–724 (1994).

67. Rodrigue, N., Lartillot, N., Bryant, D. & Philippe, H. Site interdependence attributed to tertiary 795 structure in amino acid sequence evolution. Gene 347, 207–217 (2005).

68. Robinson, D. M., Jones, D. T., Kishino, H., Goldman, N. & Thorne, J. L. Protein evolution with dependence among codons due to tertiary structure. Mol. Biol. Evol. 20, 1692–1704 (2003).

69. Chen, V. B. et al. MolProbity: all-atom structure validation for macromolecular crystallography. Acta Crystallogr. D Biol. Crystallogr. 66, 12–21 (2010).

70. Delano, W. The PyMOL molecular graphics system. http://www.pymol.org/ (2002).

71. Feng, L., Wang, J.-T., Jin, H., Qian, K. & Geng, J.-G. SH3KBP1-binding protein 1 prevents epidermal growth factor receptor degradation by the interruption of c-Cbl-CIN85 complex. Cell Biochem. Funct. 29, 589–596 (2011).

72. Kasahara, K. et al. Ubiquitin-proteasome system controls ciliogenesis at the initial step of axoneme extension. Nat. Commun. 5, 5081 (2014).

73. Lo, S.-C., Li, X., Henzl, M. T., Beamer, L. J. & Hannink, M. Structure of the Keap1:Nrf2 807 interface provides mechanistic insight into Nrf2 signaling. EMBO J. 25, 3605–3617 (2006).

74. Baker, N. A., Sept, D., Joseph, S., Holst, M. J. & McCammon, J. A. Electrostatics of nanosystems: application to microtubules and the ribosome. Proc. Natl. Acad. Sci. U.S.A. 98, 10037–10041 (2001).

75. Dolinsky, T. J., Nielsen, J. E., McCammon, J. A. & Baker, N. A. PDB2PQR: an automated pipeline for the setup of Poisson-Boltzmann electrostatics calculations. Nucleic Acids Res. 32, W665–W667 (2004).

76. Pettersen, E. F. et al. UCSF Chimera—A visualization system for exploratory research and analysis. J. Comput. Chem. 25, 1605–1612 (2004).

77. Kabsch, W. & Sander, C. Dictionary of protein secondary structure: pattern recognition of hydrogen-bonded and geometrical features. Biopolymers 22, 2577–2637 (1983).

78. Tien, M. Z., Meyer, A. G., Sydykova, D. K., Spielman, S. J. & Wilke, C. O. Maximum allowed solvent accessibilites of residues in proteins. PLoS One 8, e80635 (2013).

79. Sydykova, D. K., Jack, B. R., Spielman, S. J. & Wilke, C. O. Measuring evolutionary rates of proteins in a structural context. F1000Res. 6, 1845 (2017).

80. de Laurent, Z. R. et al. Polymorphisms in the K13 gene in Plasmodium falciparum from different malaria transmission areas of Kenya. Am. J. Trop. Med. Hyg. 98, 1360–1366 (2018).

81. Kamau, E. et al. K13-propeller polymorphisms in Plasmodium falciparum parasites from Sub-Saharan Africa. J. Infect. Dis. 211, 1352–1355 (2015).

